# Serotonergic development of active sensing

**DOI:** 10.1101/762534

**Authors:** Alireza Azarfar, Yiping Zhang, Artoghrul Alishbayli, Dirk Schubert, Judith R. Homberg, Tansu Celikel

## Abstract

Active sensing requires adaptive motor (positional) control of sensory organs based on contextual, sensory and task requirements, and develops postnatally after the maturation of intracortical circuits. Alterations in sensorimotor network connectivity during this period are likely to impact sensorimotor computation also in adulthood. Serotonin is among the cardinal developmental regulators of network formation, thus changing the serotonergic drive might have consequences for the emergence and maturation of sensorimotor control. Here we tested this hypothesis on an object localization task by quantifying the motor control dynamics of whiskers during tactile navigation. The results showed that sustained alterations in serotonergic signaling in serotonin transporter knockout rats, or the transient pharmacological inactivation of the transporter during early postnatal development, impairs the emergence of adaptive motor control of whisker position based on recent sensory information. A direct outcome of this altered motor control is that the mechanical force transmitted to whisker follicles upon contact is reduced, suggesting that increased excitability observed upon altered serotonergic signaling is not due to increased synaptic drive originating from the periphery upon whisker contact. These results argue that postnatal development of adaptive motor control requires intact serotonergic signaling and that even its transient dysregulation during early postnatal development causes lasting sensorimotor impairments in adulthood.

## Introduction

During active sensing, the position of sensory organs is adaptively controlled based on sensory and contextual information as well as learned task requirements (Nelson and MacIver, 2006; Celikel and Sakmann, 2007; Morillon et al., 2015; Dominiak et al., 2019). In the whisker system, for example, the pattern of whisking during object localization is modulated by whisker touch as animals protract their whiskers towards the presumed location of the tactile target of interest (Voigts et al., 2015). Modulation of motor (whisking) pattern by sensory input allows compensation for the change in body position in respect to the target (Carvell and Simons, 1990; Mitchinson et al., 2007; Towal and Hartmann, 2008; Voigts et al., 2008; Grant et al., 2012b). A recent study (Azarfar and Celikel, 2019) showed that this adaptive motor control develops postnatally after the 3rd postnatal week, but the developmental mechanisms of adaptive sensorimotor control are yet to be unraveled.

Postnatal development of sensorimotor circuits relies on the combination of intrinsic molecular cues and neuronal activity that is coherent across populations of neurons; while intrinsic cues ensure orderly development of the sensorimotor circuits, activity-dependent processes fine-tune the circuits thereafter (Wiesel and Hubel, 1963; Hensch, 2005; Kanold and Shatz, 2006; Erzurumlu and Gaspar, 2012; Levelt and Hübener, 2012; Martens et al., 2015). Emerging evidence suggests that neuromodulator neurotransmitters, in particular serotonin (Schubert et al., 2015), powerfully regulate the development of sensorimotor circuits. Altered serotonergic signaling during early postnatal development impairs the structure and function of neural circuits throughout the brain (Shemer et al., 1991; Whitaker-Azmitia, 1991; Moses-Kolko et al., 2005; Deiró et al., 2006; Pang et al., 2011; Kiryanova et al., 2013; Kroeze et al., 2016), results in anxiety-and depression-like behavior (Olivier et al., 2008; Kiryanova et al., 2013; Kroeze et al., 2016), hinders goal-directed navigation, and impairs object recognition (Kiryanova et al., 2013; Kroeze et al., 2016). Increasing extracellular serotonin availability during development impairs exploration well into adolescence (Kiryanova et al., 2013) while administration of select SSRIs (e.g. Fluoxetine, Sertraline, citalopram) during a critical period delay development of several reflexes and muscle strength (Deiró et al., 2006; Bairy et al., 2007; Zimmerberg and Germeyan, 2015), motor coordination (Ansorge et al., 2004, 2008; Lee and Lee, 2012) and exploration (Rodriguez-Porcel et al., 2011; Simpson et al., 2011; Miceli et al., 2017). These findings argue that changes in the serotonergic drive early in life might be critically involved in the maturation of sensory and motor circuits, which might have long-lasting consequences in adulthood.

In agreement with these observations, we previously showed that serotonin transporter knockout (SERT−/−) rats display behavioral and neuronal sensory hyperexcitability in the whisker system (Miceli et al., 2017). At the functional level, a major consequence of network excitability is the faster integration of sensory information during whisker based tactile navigation (Miceli et al., 2017). Because the barrel cortex and the primary motor cortex have reciprocal monosynaptic communication (Kleinfeld et al., 1999; Bosman et al., 2011), altered serotonergic signaling could potentially dysregulate whisker positional control during sensory navigation. Here we addressed this question using a publically available high-speed videography database of rats performing a tactile object localization task (Azarfar et al., 2018b). By quantitatively analyzing the motor control dynamics of whisker and body position, we show that sustained alterations of serotonergic signaling in SERT−/− rats, or the transient inactivation of the serotonin transporter during early postnatal development impairs emergence of adaptive motor control of whiskers, but not the body positional control in adulthood. Simulation of the mechanics of the whisker displacement showed that the absence of goal-oriented adaptive whisking reduces the mechanical forces at the whisker follicle when whiskers contact a tactile target. These results argue that previously described sensory hyperexcitability upon altered serotonergic signaling (Miceli et al., 2017) is not due to increased mechanosensory drive, but that the hyperexcitability originates in the central nervous system. In short, postnatal development of adaptive motor control requires intact serotonergic signaling, and even transient changes in serotonergic signaling during development cause long-term sensorimotor disturbances in adulthood, impair close-loop sensorimotor control of whisker position, reduce the precision in motor control during goal-directed navigation, and limit the touch induced force transmitted to the whisker follicles.

## Materials and Methods

All experimental procedures have been performed in accordance with the Dutch and European laws concerning animal welfare after the institutional ethical committee approval. The raw data have been previously made publically available as a part of the tactile navigation database (Azarfar et al., 2018b). It is freely distributed under a Creative Commons CC-BY license which permits unrestricted use, distribution, and reproduction of the data in any medium provided that the database (Azarfar et al., 2018b) is properly cited.

The experiments were performed using 38 male Wistar rats as described before (Azarfar et al., 2018b). In short, animals shuttled between two elevated platforms with a variable gap-distance in between them as they searched for a tactile target in their immediate environment in the absence of any visible light. Animal behavior on this so-called spontaneous gap crossing task (Celikel and Sakmann, 2007; Voigts et al., 2008, 2015) was recorded by a robotic high-speed camera using infrared back-lights. Videos were analyzed using custom-written routines in Matlab to extract whisking and body locomotion dynamics at a sampling resolution of 5 ms.

### Animals

Animals were randomly assigned to two groups: Half of the rats were genetically modified or pharmacologically treated to alter serotonergic neurotransmission. Serotonin transporter knockout rats (Slc6a41Hubr, SERT−/−, N=14) were originally generated on a Wistar background by N-ethyl-N-nitrosurea (ENU)-induced mutagenesis (Homberg et al., 2007). To transiently interfere with the serotonergic signaling after birth, fluoxetine hydrochloride (10 mg/kg/day, Sigma Aldrich), a selective serotonin reuptake inhibitor, was dissolved in water and administered orally to wild type Wistar rats (N=5) for 7 days after P1. The other half of the rats were wild type (N=14) and vehicle (tap water) controls (N=5).

### Analysis of tactile exploration statistics

All recorded images were analyzed off-line in Matlab. A background image, acquired before the subject’s entrance into the field of view, was subtracted from all other frames to aid with moving object detection. This background image was also used to identify the stationary tactile target, i.e. the elevated platform animals are searching for, by finding the transition point from low to high brightness in the illumination.

For the analysis of the head (body) and whisker position, the head contour was extracted using a standard contrast based edge detection approach. The positions of ears and nose were used to triangulate the head direction. The angular deviation of the nose from the head center plane (between the ears) allowed calculation of the head angle with respect to the tactile target and the relative nose distance to target. A subset of whiskers (N=6 whiskers/trial, N=3 whiskers/snout) were tracked manually; all whisker contacts with the target by micro and macro vibrissae were marked manually. Animals had all of their whiskers intact throughout the experiments. The angular displacement of the whisker base in respect to target was computed for each tracked whisker. Only data from whisker distances (i.e gap distance 14.5<x<~18 cm), where animals collected tactile information from the target using solely their whiskers prior to the decision-making, were analyzed. Thus tactile exploration using the touch receptors embedded in the skin (as it invariably happens at short (nose) distances (Celikel and Sakmann, 2007)) do not interfere with the sensorimotor behavior (Celikel and Sakmann, 2007) and the adaptive sensorimotor control of whisker position during tactile navigation (Voigts et al., 2008, 2015).

The whisker position in respect to the target platform was calculated as the Euclidean distance between them as per previous studies (Voigts et al., 2008, 2015; Miceli et al., 2017). Angular protraction of the whisker was quantified at the whisker base, with respect to the position of the whisker at rest, i.e. the midpoint between the protraction and retraction set-points in a whisk cycle (Figure 1A). Whisker tip position and relative position of the tip of the nose with respect to the tactile target were extracted using custom routines. The absolute distance between the two platforms was defined as the target distance (in respect to the home platform).

**Figure 1.**
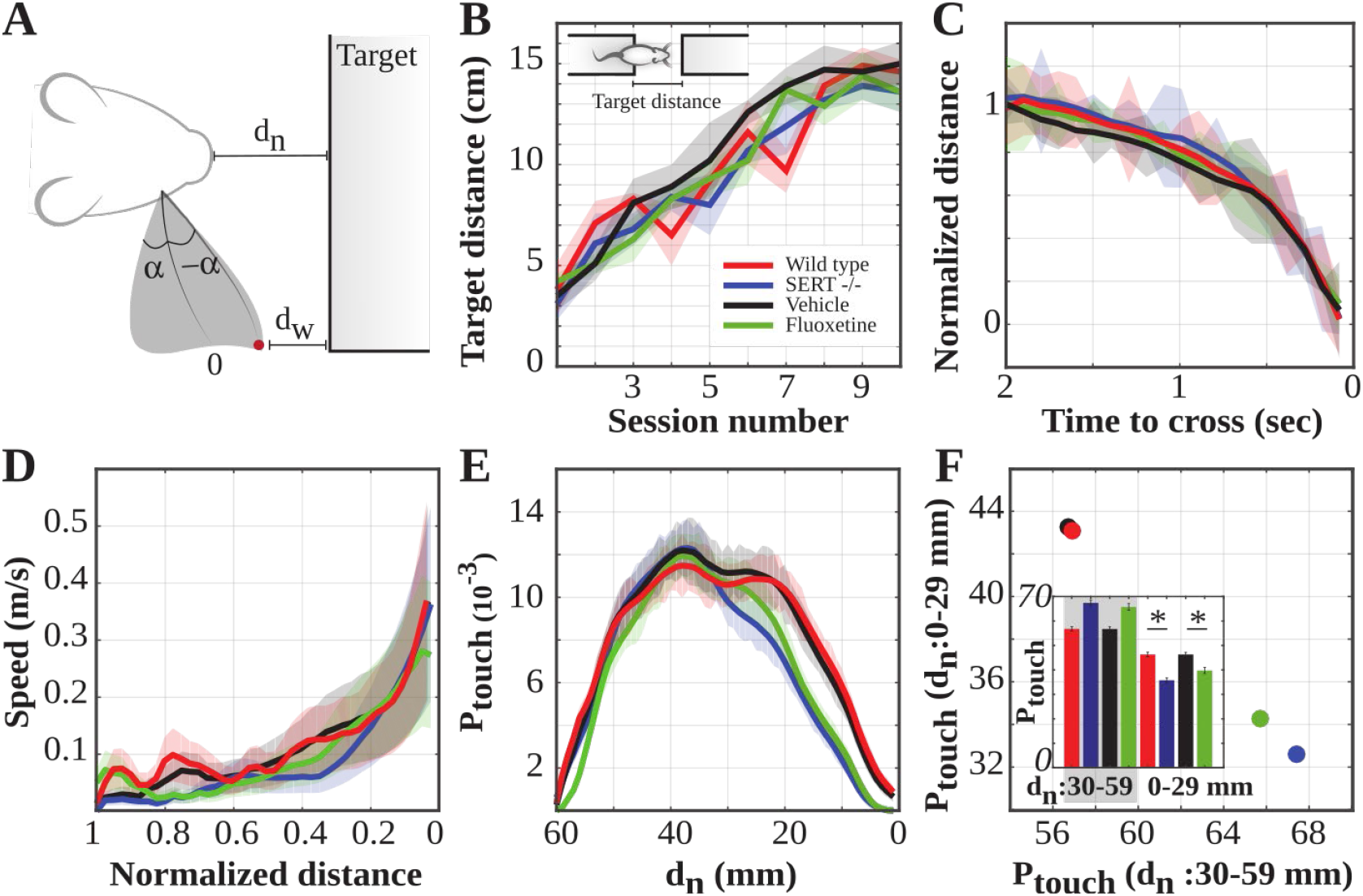
The role of serotonin in goal-directed sensorimotor navigation. **(A)** Behavioral parameters of interest. The red dot denotes the whisker tip; α is the angular displacement from the mid-point between the most retracted (i.e. retraction set-point, α) and protracted positions (protraction set-point, - α) of the whisker during whisking. dn is the relative nose distance to target and d_w_ symbolizes the tracked whisker’s tip distance to target. **(B)** Learning curves for the four groups (Group effect: F=1.83; p=0.141, 2-way ANOVA with df=3). **(C)** Normalized distance from the target during the last 2 seconds of the exploration task (Group effect: F=0.835; p=0.475, 2-way ANOVA with df =3). Note that the average tactile exploration duration was less than a second. **(D)** Speed of locomotion during the last one second of the object localization task (Group effect: F=0.670; p=0.517, 2-way ANOVA with df=3). **(E)** Normalized probability of contact with the target platform as a function of the relative distance to the target (Group effect: F=22.931; p <0.001, 2-way ANOVA with df =3). **(F)** Normalized probability of touch based on the relative position of the animal to the target. Animals with altered transient (fluoxetine) and sustained (SERT−/−) serotonergic signaling required less number of contacts with the target platforms prior to successful object localization (Vehicle vs Fluoxetine, p<0.05; Wild type vs SERT−/−, p<0.05; t-test). Data are presented as mean and the standard deviation from the mean, unless stated otherwise.

### Mechanical modeling of whisker bending

Whiskers can be modeled as a flexible beam that transmits mechanical information to mechanoreceptors in the follicle. Whisker geometry, density, and its elastic moduli govern the mechanical information at the follicle. This mechanical “simplicity” allows mathematical approximation of the tactile (mechanical) inputs transmitted by the vibrissae during active tactile exploration.

Whisker shape can be approximated by a quadratic fit (a parabola) (Towal et al., 2011). Given that the density of the keratinized tissue diminishes almost linearly from base to tip (Williams and Kramer, 2010; Hires et al., 2013b), the whisker’s elastic moduli govern its bending stiffness. The two most relevant moduli to whisker mechanics are Young’s modulus (E) and shear modulus (G). Young’s modulus is essential to model the whisker in 2D, as shear modulus is significant only when a force is applied to the whisker surface while the opposite side of the whisker is held constant by another equal force. The equation below translates the curvature (k) at each point (s) along the whisker to bending moment (M), given the stiffness of the whisker at each point. This bending moment (the product of stiffness and the curvature), is used to estimate the signals that rats experience during whisker deflection (Solomon and Hartmann, 2006; Boubenec et al., 2012; Quist and Hartmann, 2012; Hires et al., 2013a; Pammer et al., 2013).

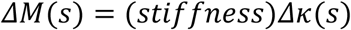

Whisker contact induced mechanical information at the whisker follicle was approximated by calculating the bending moment (M), the axial force (F_x_,which is a force directed along the length of the whisker near the base), and the transverse force (F_y_) that is perpendicular to the axial force (Fig.4A) as described previously using Elastica 2D in MATLAB (Solomon and Hartmann, 2006; Quist and Hartmann, 2012). The body locomotion towards the object was based on the experimental observations and reflected the approach trajectory quantified above. Under these constraints body-to-target position along with the egocentric whisker position (alpha of the parabola fit to the whisker) govern where along the whisker shank contact with the target would happen, therefore determining the force dynamics at the whisker base.

## Results

Serotonergic signaling is critically involved in the development and maturation of circuits throughout the brain, however, its role in the emergence of closed-loop sensorimotor computations for adaptive motor control is unknown. Here, we addressed this question on a tactile object-localization task with genetically engineered and pharmacologically treated rats and computational simulations.

### Task performance and development of goal-oriented motor control strategies

Sensory circuits represent the world in egocentric coordinates but body motion and moving appendages (e.g. whiskers, limbs, and head) necessitate discounting the changes in body position for the integration of sensory information in space and time. In the adult whisker system, this is achieved by adapting the whisker protraction angle to the change in body position with respect to the tactile target to perform a form of vector computation where whiskers and body are treated as coupled manifolds (Voigts et al., 2015). This computation is learned after the third postnatal week upon sequential emergence of adaptive motor control of body and whiskers (Grant et al., 2012a; Azarfar and Celikel, 2019). If serotonergic signaling is required for the adaptive sensorimotor computation, body or whisker motion control is expected to be impaired by altered serotonergic signaling during development.

To observe the motor control dynamics during or following altered serotonergic signalling, four groups of rats were studied in the gap-crossing task (Celikel and Sakmann, 2007; Voigts et al., 2008, 2015; Pang et al., 2011; Juczewski et al., 2016; Miceli et al., 2017): serotonin transporter knockout (SERT−/−) rats had constitutive deletion of the transporter, while rats postnatally exposed to Fluoxetine and experiencing transient pharmacological inactivation of the transporter during development allowed the assessment of serotonergic contribution to the emergence of sensorimotor control. Wild-type controls (SERT+/+) and vehicle-treated groups (Vehicle) served as controls. All behavioral observations were performed in adulthood (~ postnatal day (P) 65).

#### All groups successfully acquired the task

Animals increased the maximum gap-distance at which they successfully located the tactile target by four-folds over the course of 10 training sessions (Fig.1B). In every session animals were tested at multiple gap-distances (range: 3-18 cm), each constituting a trial. After the completion of a trial, gap-distance was randomly selected from a normal distribution where the distribution mean increased and the variance was reduced with the increasing number of training sessions completed on the task (Miceli et al., 2017; Azarfar et al., 2018b).

#### All groups employed a similar locomotor strategy to accomplish the task

Animals approached the target at a comparable rate (Fig.1C) and speed (Fig.1D). Despite the similarity in approach trajectories across the four groups, animals that received transient (with Fluoxetine exposure) or permanent SERT deletion (SERT−/−) made significantly less number of contacts with the target during the last phase of the approach (i.e last 29 mm; Figure 1E, 1F) as previously shown in juvenile SERT−/− rats (Miceli et al., 2017). Since SERT deletion does not change animals’ basic body locomotion strategies, the reduction in sensory exploration might be due to distinct whisking strategies employed after/during serotonergic dysregulation.

### Adaptive sensorimotor control of whisker position

During tactile exploration of stationary objects animals protract their whiskers to the position in space which they recall as the location of the tactile target (Voigts et al., 2015). In agreement with these observations, we found that as animals approached a stationary target, the relative distance between the target and their whisker tip position was reduced independent from whether animals received acute or chronic intervention with serotonergic signalling (Fig.2A).

**Figure 2.**
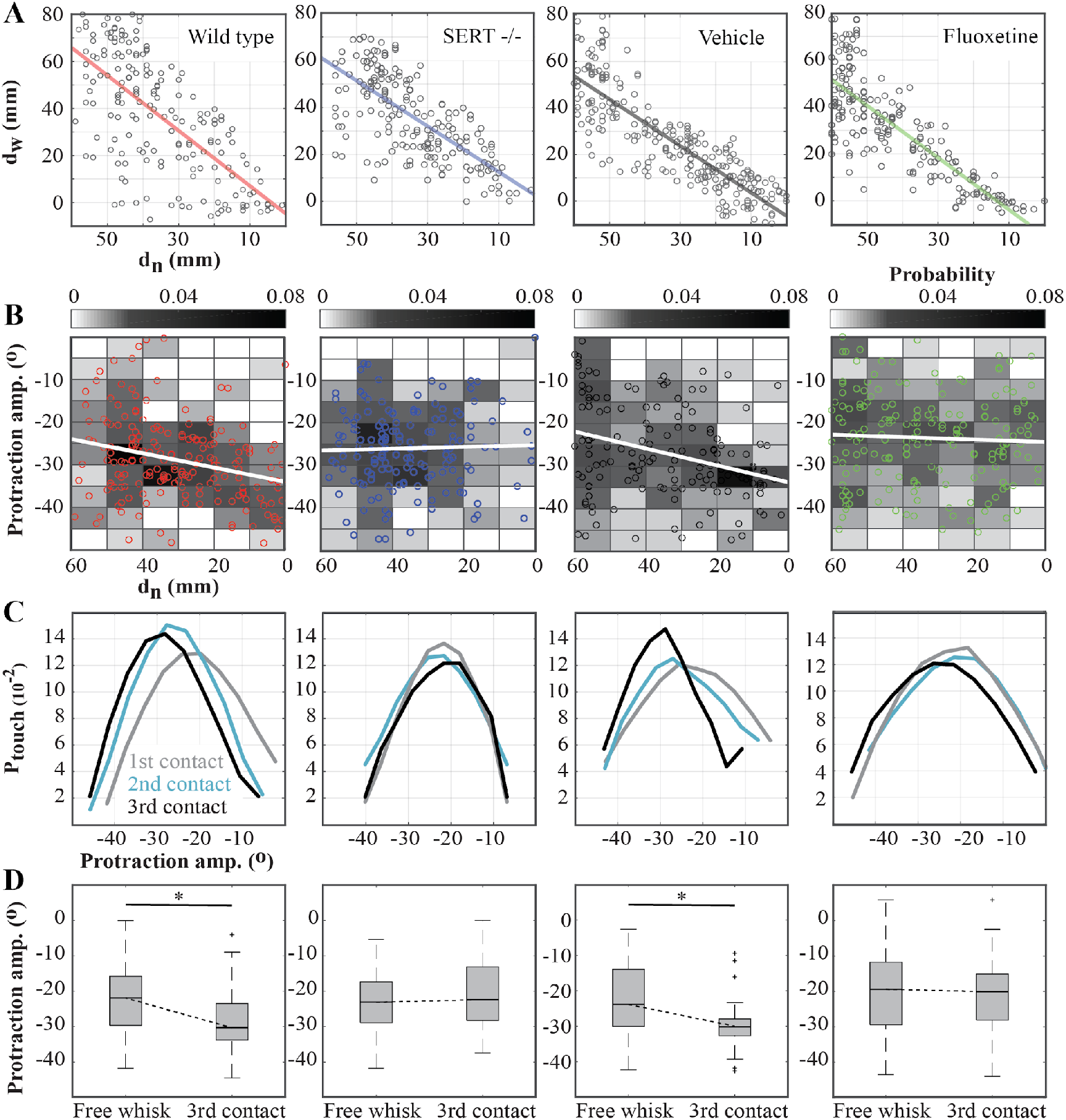
Serotonergic regulation of adaptive sensorimotor control of whisker protraction. **(A)** Whisker tip distance to target (d_w_) with respect to the relative distance of nose to target (d_n_). From left to right: Wild-type (r^2^=.47, Nose distance effect: F=2.92; p<0.001), SERT−/− (r^2^=.50, Nose distance effect: F=5.42; p<0.001), vehicle (r^2^=.75, Nose distance effect: F=9.29; p<0.001) and fluoxetine (r^2^=.72, Nose distance effect: F=7.58; p<0.001). **(B)** Most protracted angle at each whisk cycle with respect to the relative distance to target. From left to right: Wild-type (Fit slope: −0.17 deg/mm, Nose distance effect (ANOVA): F=10.59; p<0.01), SERT−/− (Fit slope: 0.02 deg/mm, F=0.925; p=0.061), vehicle (Fit slope: −0.22 deg/mm, F=47.46; p<0.01) and fluoxetine (Fit slope: 0.02 deg/mm, F=1.649; p=0.017). The 2D histogram of the data is represented at the background and a representative sample data from a random single trial is plotted on top. **(C)** Normalized histograms of protraction angles binned across consecutive whisker contacts with the target (Touch sequence effect (2-way ANOVA) -- Wild-type: F=16.84, p<0.001 with df=2; SERT−/−: F=1.46, p=0.098 with df =2; vehicle: F=7.98; p<0.001, df=2, Fluoxetine: F=2.99, p=0.053 df=2). **(D)** Change in whisker protraction angle between free-whisking and the third whisker contact with target (ANOVA, Wild-type touch sequence effect: F=10.126; p<0.001 with df=1, SERT−/−: F=2.13; p=0.147 with df=1, vehicle: F=26.369; p<0.001 with df =1, Fluoxetine: F=1.55; p=0.214 with df=1).

The adaptive control of whisker position is a function of the coupling between two moving manifolds, i.e. body movement and whisker motion. It was suggested that the functional contribution of the adaptive changes in whisker protraction might be to account for the changes in body position (Voigts et al., 2015). If the brain were to perform vector computation to calculate the relative distance of the target by considering the displacement of the body and the whisker position, reduction of the protraction angle would allow to minimize the error in stimulus location estimates. The observation that for all groups whisker’s tip position converges to the expected target position (Fig.2A), however, does not imply that animals adaptively control the whisker protraction. This is because the whisker tip position is also a product of the body position across whisk cycles. Therefore, we next quantified the change in whisker protraction as animals approached to the target.

As it was shown previously, adult (Voigts et al., 2008, 2015), but not juvenile (Azarfar and Celikel, 2019), animals utilize adaptive sensorimotor control by increasing the maximum whisker protraction angle while decreasing whisk amplitude as they get closer to the target. This protraction angle maximization is mediated by advancing the midpoint around which the whisker swings at each cycle (Voigts et al., 2015). Animals with altered serotonergic signaling, but not their corresponding controls, lack this adaptive sensorimotor control independent from whether the serotonergic dysfunction was transient (i.e. fluoxetine group) or persistent (SERT−/−) (Fig.2B). Sensory information required for the adaptive motor control is acquired over the last ~3 whisk cycles as animals infer the target location based on this “prior” to control whisker position in the next whisk cycle (Voigts et al., 2015). The progressive adaptation of the whisker protraction angle upon successive contacts with the target (Fig.2C) indicates that the control groups perform motor planning based on the recent sensory information. By integrating the sensory knowledge iteratively by the third touch, they increase the protraction angle, extend their whiskers towards the target (Fig.2D). In SERT−/− and fluoxetine groups, however, the whisker protraction angle is independent of the prior whisker contacts (Fig.2C,D). These results show that even transient changes in serotonergic signaling during development have long-lasting consequences for adaptive sensorimotor control in adulthood.

#### Mechanical forces transmitted to the whisker follicle upon whisker contacts are reduced upon serotonergic dysregulation

Whiskers are specialized sensory hairs. As such, they do not contain any sensory receptors along their shanks; the mechanical displacement of the whisker is transmitted to the whisker follicle for sensory transduction (Rice et al., 1986). During non-adaptive whisking (as observed in SERT−/− and fluoxetine animals) the changes in follicle displacement upon whisker contact are solely due to a change in body position (e.g. distance, head angle) with respect to the target. During adaptive whisking (as performed by wild type and control animals), the changes in the whisking pattern also contribute to the mechanical forces at the follicle. Thus the quantification of the force in the follicle across non-adaptive and adaptive whisking could help to unravel the sensory contribution of adaptive motor control.

To quantify the force impinging on the follicle we implemented a mechanical model of the rat whisker (see Methods for details). Whisker displacement during active palpation onto a stationary target was simulated as animals gradually approached the target according to behaviorally observed trajectories (Fig.1). In all *in silico* experiments, only the last 3 cm of the approach was simulated where animals with serotonergic dysregulation contacted the target less often than their counterparts in the corresponding control groups (Fig.1E-F). For the simulation of non-adaptive whisking, the maximum protraction angle of the whisker was kept constant (~ 27°). During adaptive whisking the protraction angle gradually increased from 27° to 37° along the locomotion path per experimental observation (Fig.2). The results showed that mechanical forces transmitted to the whisker follicle upon whisker contact is increased as the animal approached the target (see red traces, Fig.3) independent from the whisking strategy employed, due to the reduction in the relative distance between the target and the body. Lack of adaptive whisking resulted in systematic reduction of the force transmitted to the whisker along three axis (blue traces, Fig.3). These results argue that adaptive whisking increases the force transmitted to the whisker base, increasing the feed-forward excitatory drive upon whisker contact. Lack of adaptive whisking after fluoxetine treatment or SERT−/− reduces forces in the whisker base upon whisker contact (Fig.3), thus reducing the sensory information originating from the periphery.

**Figure 3.**
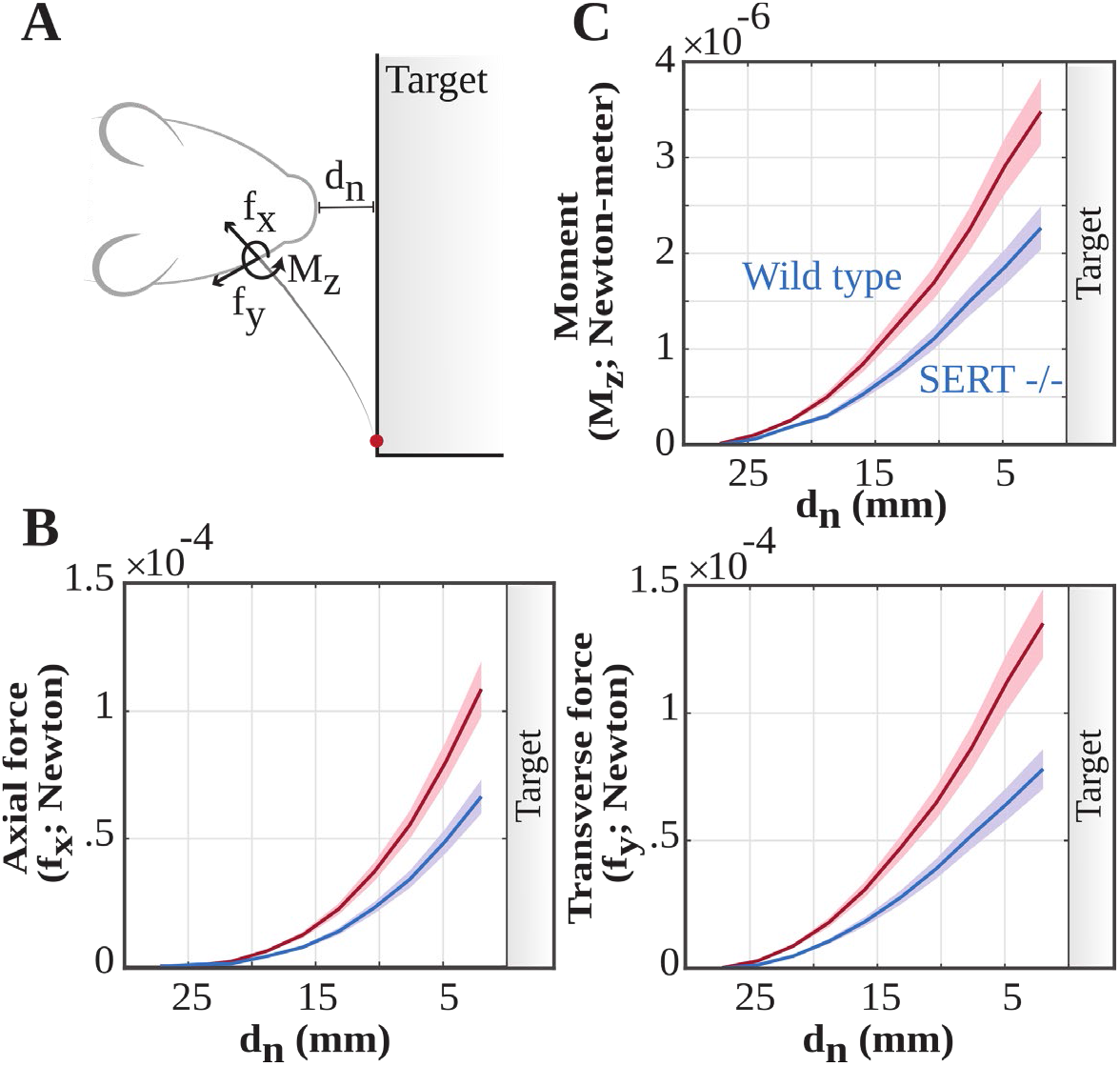
Lack of adaptive sensorimotor control upon altered serotonergic signaling reduces the force transmitted along the whisker upon contact with a tactile target. **(A)** Whisker contacts with objects in the plane of whisking change the axial force (F_x_), transverse force (F_y_) and the reaction moment (M_z_) at the whisker base, and leads to mechanoreceptor activation to initiate bottom-up propagation of the sensory information. **(B)** Mechanical forces at the whisker base upon whisker contacts as the animal approached the target (Axial force genotype effect: F=88; p <0.001, 2-way ANOVA with df=1, Transverse force genotype effect: F=113; p <0.001, 2-way ANOVA with df =1). **(C)** Change in Moment (M_z_) at the follicle upon whisker contact (Genotype effect: F=56; p <0.001, 2-way ANOVA with df=1). The data presented as mean ∓ std, blue denotes non-adaptive whisking conditions (SERT−/−, fluoxetine) and red adaptive whisking conditions (wild type, vehicle).

### A network model of adaptive whisking

Sensory exploration and motor control are coupled processes. Considering that lack of adaptive whisking reduces mechanical forces traveling along whiskers upon whisker touch (Fig.3), this change in the sensory drive could potentially alter the motor control of whisker position in subsequent whisk cycles. To address this question without the confounding variables (including compensatory change in body position) of navigation in freely behaving animals, we used a simplified graph based network model of whisking (Circuits of whisking; github.com/DepartmentofNeurophysiology/Circuits-of-whisking).

A computational circuit that could perform adaptive sensorimotor control necessarily requires information from sensory circuits about the stimulus availability as well as motor control circuits that perform phase to motor signal transformation given the current state of the sensory information. Based on the known coding properties of the neurons along sensorimotor circuits, and the connectivity between them (see Discussion), the graph network consists of the following nodes (Fig.4A): 1) primary somatosensory cortex (S1; barrel cortex) where stimulus properties are encoded (Brecht and Sakmann, 2002; Ganguly and Kleinfeld, 2004; Crochet and Petersen, 2006; Curtis and Kleinfeld, 2009; de Kock and Sakmann, 2009; Lundstrom et al., 2010; Azarfar et al., 2018a); 2) primary motor cortex (M1) which provides adaptive motor control for whisker protraction (Berg and Kleinfeld, 2003a; Brecht et al., 2004; Diamond et al., 2008; Petersen, 2014; Sreenivasan et al., 2016), through recursively adjusting the amplitude and midpoint of whisking envelope (Hill et al., 2011); 3) central pattern generators (CPGs) that control phasic motion of whiskers (Gao et al., 2001; Cramer and Keller, 2006; Kleinfeld et al., 2015); 4) superior colliculus (SC) which translates phase and amplitude information to motor control commands for facial motor nucleus (FMN) to drive whisking (Hemelt and Keller, 2008); 5) dorsal raphe nucleus (DRN) that regulates excitability in cortical and subcortical (sensorimotor) nuclei (Schubert et al., 2015); and 6) a control circuit, plausibly the barrel cortex (Matyas et al., 2010), that triggers whisker retraction upon stimulation to maintain touch duration (Azarfar and Celikel, 2019). In this model output of each node is a transfer function rather than a time and/or rate varying action potentials. Please note that the aim of this model is not to mechanistically explain how the brain performs sensorimotor computation. Rather, it is to provide the minimal circuit requirements for adaptive control of whisker position (see Discussion).

**Figure 4.**
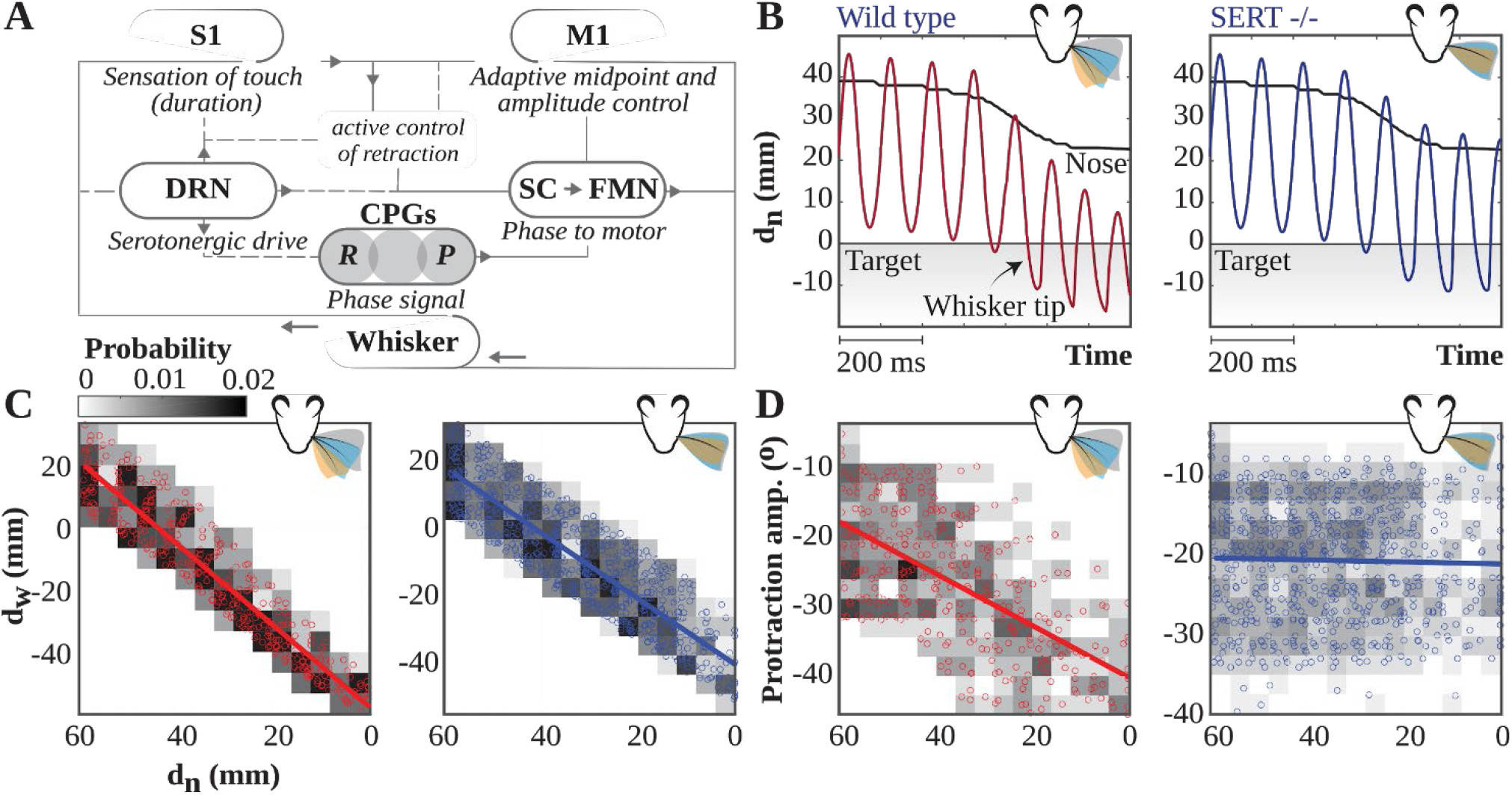
A computational circuit model of adaptive sensorimotor control. **(A)** Circuit components in the network: Barrel cortex subregion of the primary somatosensory cortex (S1), vibrissal motor cortex (M1), dorsal raphe nuclei (DRN), Superior colliculus (SC), central pattern generator (CPG) and facial motor nuclei (FMN) are modeled. See Discussion for details on the known anatomical and functional projections along this network. **(B)** Relative distance of a whisker tip to the tactile target during simulated adaptive whisking (red; left figurines) and simulated non-adaptive whisking (blue; right). The black line is an experimentally observed, randomly selected approach trajectory in a 2D plane, i.e. change in relative Euclidean distance to the target (d_n_) as a freely behaving animal approaches a stationary tactile object. In adaptive whisking, sensory information modulates the motor command: protraction angle increases given the sensory information collected prior to the current whisk cycle, whisking amplitude decreases, and whisker retraction is actively controlled to keep the touch duration constant. Figures summarize the experimental observations regarding whisker protraction amplitude, set-point and mid-point in the respective group of animals. **(C)** Simulated tip distance to target (d_W_) in relation to body-to-target position. Left: adaptive whisking (red, r^2^=0.89); right: non-adaptive whisking (blue, r^2^=0.78). **(D)** Protraction amplitude, in respect to mid-point, versus the nose distance d_n_ from target *in silico*. Left: adaptive whisking (red, r^2^=0.40); right: non-adaptive whisking (blue, r^2^<0.01).

Simulations in this circuit showed that adaptive whisker protraction (Fig.4B, left) is an emergent computation and can be dysregulated by either removal of the serotonergic reuptake or increasing the excitability in the sensory cortex, which was previously shown in SERT−/− animals (Miceli et al., 2017) (Fig.4B, right). Although simulated animals, similar to rats (Fig.2), continue remapping whisker position as they approach the tactile target (Fig.4C), in circuits simulations without adaptive sensorimotor control do not result in a change in increased whisker protraction, similar to the observations in SERT−/− and fluoxetine animals (Fig.4D, compare it to Fig.2B).

#### Adaptive whisking improves tactile scanning resolution and this difference in whisking pattern is sufficient to explain the reduced sensory exploration during serotonergic dysfunction (seen in Fig. 1)

The aforementioned computational model might help to unravel whether the reduced likelihood of tactile exploration observed in the SERT−/− and Fluoxetine groups is a product of the lack of adaptive motor control. To address this question we simulated sensorimotor exploration of a stationary target *in silico* (Fig.5). In this experiment the object was “touch transparent”, as such contact with the target did not change whiskers’ motion trajectory. This is akin to the “virtual whisker tip position” mapping described previously (Voigts et al., 2015) and ensures that intended whisker tip position could be visualized (Fig.5A).

**Figure 5.**
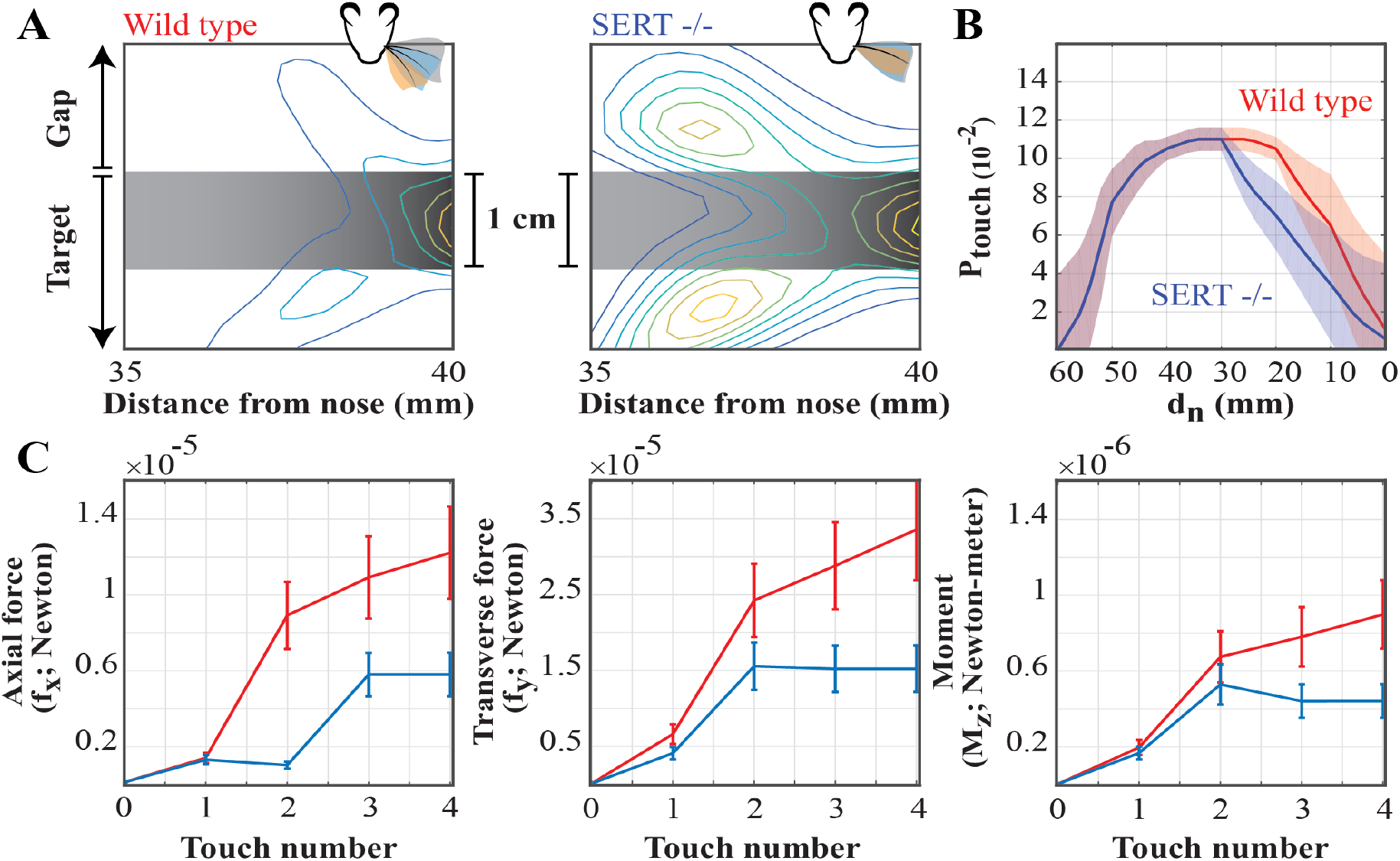
Lack of adaptive motor control alone is sufficient to explain the sensory exploration pattern upon altered serotonergic transmission. **(A)** Whisker tip position during exploration of a stationary target *in silico*. Grey shaded area represents the edge of tactile target. Density plots quantify the relative position of the whisker tip during tactile exploration with (Wild type: Contours are drawn with a step size of 16.5%, range: 16.5-100%) and in the absence of (SERT−/− contours: step size of 12%, range: 9-100%) adaptive control of whisker protraction. **(B)** Probability of whisker contact with target *in silico*. Red: Adaptive whisker protraction (as seen in rats in wild type and vehicle injected groups); Blue: Non-adaptive whisker protraction (as performed by SERT−/− and after transient pharmacological intervention. **(C)** Mechanical forces (F_x_ and F_y_) and Momentum (M_z_) evolution at the whisker base (mean ∓ std) during simulated whisker contacts with tactile target *in silico*. Color code as in B. The approach trajectory for the adaptive and non-adaptive whisker protraction are based on behavioral observations. Touch number 0 in the X axis refers to the last whisking cycle prior to first contact with the target.

The simulations showed that tactile navigation using adaptive whisker protraction results in *in silico* whiskers being positioned at the virtual target (Fig.5A, left) while non-adaptive whisking introduce localization errors (Fig.5A, right). These results could potentially explain the reduced tactile exploration observed in experiments (Fig.1E) with SERT−/− and fluoxetine injected animals as the likelihood of whisker contacts with the target observed *in silico* closely resemble the experimental observations (compare Fig.1E to Fig.5B). After the first touch event, the whisker motor commands are modulated and directed towards the target in adaptive whisking. This target alignment directly increases the probability of touch events.

Considering that the body locomotion together with the whisking pattern govern the forces at the follicle, the localization errors observed during non-adaptive whisking could contribute to the reduction in mechanical forces transmitted to the follicle upon whisker contacts. Simulations showed that the contact induced forces were indeed significantly smaller during non-adaptive whisker protraction (Fig.5C). Larger touch induced mechanical information (i.e. force) at the follicle, improved tactile resolution (reduced localization error), and increased likelihood of sensory exploration argue that adaptive whisking results in higher signal to noise ratio during sensory acquisition.

## Discussion

Here we showed that persistent or transient SERT inactivation induce lasting impairment of sensorimotor computation, and interfere with the development of adaptive sensorimotor control during tactile object localization. Specifically, after serotonergic interventions animals fail to integrate sensory information to regulate the whisker positions in the subsequent whisk cycles (Fig.2B). Nonetheless these rats are able to perform object localization successfully, similar to the wild-type and vehicle groups. To address whether changes in sensorimotor strategies alter sensory information mechanically transmitted along the whisker upon whisker contact, we modeled the forces generated at the follicle during active whisking based on the behavioral data. The results showed that adaptive whisking maximizes the forces transmitted along the whisker, although this nonlinear mechanical gain modulation is spatially constrained. A computational circuit model of adaptive versus non-adaptive (uniform) whisking showed that inactivation of the communication between primary somatosensory and primary motor cortices impairs adaptive whisking sensorimotor control. The results also indicated that adaptive whisking improves tactile scanning resolution and confirmed the finding that adaptive whisking strategy increases sensory information transmitted during tactile exploration. Higher scanning resolution and stronger signal representation at the follicle translates to higher signal to noise ratio in sensory acquisition. The outcome of the simulation further proposed that the difference in whisking pattern between the two groups is sufficient to explain the sensory exploration pattern after alterations in serotonin transmission.

### Altered excitation/inhibition balance might explain serotonergic phenotypes

Changes in the serotonergic drive early in life might have long-term behavioral consequences including depression and anxiety (McAllister et al., 2012; Francis-Oliveira et al., 2013; Kiryanova et al., 2013; Kroeze et al., 2016), deficit in circadian rhythmicity (Kiryanova et al., 2013), reduction in body weight (McAllister et al., 2012; Kroeze et al., 2016), decreased social behavior (Kiryanova et al., 2013), reduced sexual motivation (Kiryanova et al., 2013; Vieira et al., 2013; Rayen et al., 2014), (Kiryanova et al., 2013; Kroeze et al., 2016), and might alter reward processing and learning & memory (Kiryanova et al., 2013). Impaired sensorimotor integration, upon serotonergic dysregulation, might contribute to the expression of many of these phenotypes (Francis-Oliveira et al., 2013; Kroeze et al., 2016). Altered serotonergic drive might also result in miswiring of sensorimotor circuits, and thus cause sensorimotor deficits. Elevated serotonin levels during the critical period disrupts locomotion (Bairy et al., 2007; Lee et al., 2008; Lee and Lee, 2012; Kiryanova et al., 2013; Kroeze et al., 2016), decreases novel object exploration (Rodriguez-Porcel et al., 2011; Kiryanova et al., 2013; Kroeze et al., 2016) and causes delay in development of several reflexes and muscle strength (Deiró et al., 2006; Bairy et al., 2007; Zimmerberg and Germeyan, 2015; Kroeze et al., 2016) possibly via structural changes in the circuit organization (Kiryanova et al., 2013). On the other hand, the lack of adaptive sensorimotor control in adult rats with early changes in serotonergic drive is also seen in juvenile animals which have not been fully matured yet. Interestingly, it has been demonstrated that fluoxetine exposure leads to a juvenile-like state of neurons across brain regions, including sensory cortices, in adults (Umemori et al., 2018). Accordingly, the non-adaptive whisking seen in SERT−/− rats and postnatally fluoxetine exposed rats may be a by-product of increased exploratory drive compensating for the lack of adaptive whisking. Alternatively, the sensorimotor deficits might be a consequence of an altered balance between excitation and inhibition (E/I) as elevated serotonin levels during development impairs feedforward inhibition and facilitate excitatory drive in the somatosensory system (Miceli et al., 2017). Considering that altered E/I balance is critical for preventing runaway excitation, sharpening stimulus selectivity, and increasing the overall sparseness of stimulus representations, a local change in E/I balance in sensory systems might have global consequences on the perceptual, cognitive and motor deficits (Pang et al., 2011; Murray et al., 2014; Juczewski et al., 2016; Lainscsek et al., 2019), even in invertebrates (Hughes and Celikel, 2019). Given the regulatory close-loop between serotonergic drive and sensory experience (Yan et al, submitted) the impact of the transient alterations in E/I balance could be sustained in longer time scales.

### A network model of whisking

To provide a simplified circuit model of adaptive computation where serotonergic dysregulation to sensorimotor integration can be quantitatively studied, we deployed a graph based network model whisking. The model is based on the known principles of neural representations and circuit connectivity across sensorimotor nuclei (see results and below). We have repeated the behavioral experiments (Fig.1-2) *in silico* to validate our model, and finally used it to address the circuit mechanisms of adaptive whisking.

Behavioral and in silico experiments showed that control animals (wild-type and vehicle groups) modulate the whisker protraction based on the recent sensory information by adjusting the midpoint of whisking. In control animals peak to peak amplitude of whisking decreases and whisking rhythm is regulated to keep the contact duration constant independent from the relative position of the body and whiskers in respect to the target. M1 is a possible candidate that controls the amplitude and midpoint of the envelope of whisking (Berg and Kleinfeld, 2003a; Brecht et al., 2004; Diamond et al., 2008). Hill et al. (Hill et al., 2011) found that the majority of single units in vM1 cortex code for variation in amplitude and midpoint of whisking. Since these motor representations are not influenced by inactivation of the trigeminal sensory input, these signals should be generated by a central source, plausibly in M1. In our whisking network model, we have a modulatory unit that applies the same controls on whisking pattern upon activation (see Figure 4A). When simulated whiskers contact the target, incoming sensory information drives the M1 module to apply goal-oriented modulation on whisking pattern. The peak to peak amplitude decreases and the maximum protraction angle increases.

Touch events modulate whisking by driving the adaptive motor control as well as through a regulatory circuit that keeps the duration (duty cycle) constant (Azarfar and Celikel, 2019). S1 (barrel cortical) neurons encode the touch event and its duration both at the single-cell and population levels (Brecht and Sakmann, 2002; O’Connor et al., 2002; Ganguly and Kleinfeld, 2004; Crochet and Petersen, 2006; Derdikman et al., 2006; Ferezou et al., 2006; Hentschke et al., 2006; Yu et al., 2019). S1 spiking correlates by rapidly varying signals that represent the phase of the motion during rhythmic whisking (Crochet and Petersen, 2006; Curtis and Kleinfeld, 2009; de Kock and Sakmann, 2009; Lundstrom et al., 2010). S1 is also linked to adaptive whisking; whisking increases phase-locking between vibrissa movement and electrical activity in barrel cortex (Brecht and Sakmann, 2002; O’Connor et al., 2002; Ganguly and Kleinfeld, 2004; Crochet and Petersen, 2006; Derdikman et al., 2006; Ferezou et al., 2006; Hentschke et al., 2006) while targeted stimulation of S1 results in whisker retraction (Matyas et al., 2010). In our in silico model, the S1 module detects the contact timing and whisker phase during a touch event. Information from S1 is ultimately integrated with the reafference copy (Crapse and Sommer, 2008) of the control signal in M1 to calculate the error between the planned whisking path and the current location (i.e. interrupted path) upon a contact event. Touch duration in S1 is further used to retract the whisker which ensures the constancy of touch duration. This calculation could emerge in a variety of brain regions (Matyas et al., 2010), including the posterior parietal cortex (Mohan et al., 2017).

In free whisking, rhythmic motion is the dominant mode of whisking (Berg and Kleinfeld, 2003b). However, in adaptive whisking this rhythmic movement is altered by adjusting the whisk amplitude and midpoint of whisk cycle (Mehta et al., 2007; O’Connor et al., 2010; Voigts et al., 2015). M1 could have an instructive role over this cyclical pattern of whisking (Gao et al., 2003). In our model we have two central pattern generator (CPG) modules, one for protraction, the other retraction generation. In the absence of sensory input (free whisking), the output of these two modules directly govern the whisking pattern. In case of contact, the M1 modules manipulates their output to instruct adaptive whisking, possibly via superior colliculus (Hemelt and Keller, 2008).

The serotonergic system contributes to sensorimotor system prominently through its projections via DRN (Commons, 2015; Schubert et al., 2015). Although these projections target most of the neocortex, in the current model, DRN regulates only a limited subset of neural loci as shown in Fig.4 as our focus is to provide a minimum circuit model that could drive adaptive sensorimotor control.

The final motor command (whisker angle/phase) is connected to a virtual model of whisker (Towal et al., 2011; Quist and Hartmann, 2012). Our whisker is modeled with a parabola and bends upon contact. We use the egocentric information of the whisker position combined with allocentric information of body-to-target position of the animal in the simulation as input to our whisker model. Using this model we calculate the bending along the whisker and the forces at the whisker’s base. The body locomotion (allocentric) information in simulation is learned and determined through experimental data of rats performing gap crossing task. Using this in-silico whisker model, the consequences of adaptive motor control on sensory acquisition is simulated.

### Outlook

The present study demonstrates that reduction of SERT during early development, either by blocking 5-HTT using an SSRI (fluoxetine) or by genetic deletion of SERT, have long-term effects on sensorimotor computation and impairs emergence of adaptive whisking, in agreement with the observations on the serotonergic contributions to motor development (Kroeze et al., 2016). As a result, whisker contacts transmit less mechanical information to whisker follicle. Considering our previous observations on the reduction of inhibitory drive and increased feedforward excitation in the primary somatosensory cortex (Miceli et al., 2017), and the observations that SERT−/− deletion reduces the thalamocortical projections targeting the cortical layer 4 (Miceli et al., 2013), it is tempting to speculate that the change in cortical excitability is a compensatory change to facilitate the detection of weak signals originating from the periphery. Regulating the excitability of inhibitory neurons in Layer 4 in a cell type specific manner during object localization will provide a mechanistic insight into the neural basis of touch sensation. Re-balancing excitatory and inhibitory drive in the somatosensory cortex will also alter the communication between S1 and M1, is expected to rescue the adaptive sensorimotor control even in adulthood.

